# Tubulin Regulates the Stability and Localization of STMN2 by Binding Preferentially to Its Soluble Form

**DOI:** 10.1101/2025.02.27.640326

**Authors:** Xiang Deng, Gary Bradshaw, Marian Kalocsay, Timothy Mitchison

## Abstract

Loss of the tubulin-binding protein STMN2 is implicated in amyotrophic lateral sclerosis (ALS) but how it protects neurons is not known. STMN2 is known to turn over rapidly and accumulate at axotomy sites. We confirmed fast turnover of STMN2 in U2OS cells and iPSC-derived neurons and showed that degradation occurs mainly by the ubiquitin-proteasome system. The membrane targeting N-terminal domain of STMN2 promoted fast turnover, whereas its tubulin binding stathmin-like domain (SLD) promoted stabilization. Proximity labeling and imaging showed that STMN2 localizes to trans-Golgi network membranes and that tubulin binding reduces this localization. Pull-down assays showed that tubulin prefers to bind to soluble over membrane-bound STMN2. Our data suggest that STMN2 interconverts between a soluble form that is rapidly degraded unless bound to tubulin and a membrane-bound form that does not bind tubulin. We propose that STMN2 is sequestered and stabilized by tubulin binding, while its neuroprotective function depends on an unknown molecular activity of its membrane-bound form.

## INTRODUCTION

Stathmins are small phosphoproteins that bind to soluble tubulin through a conserved “Stathmin Like Domain” (SLD). Vertebrate genomes encode five stathmins: STMN1, STMN2 (SCG10), STMN3 (SCLIP), STMN4 (RB3) and STMND1. STMND1 has a more divergent SLD compared to the others and was only recently recognized to be a tubulin-binding stathmin [Deng et al., 2024]. STMN1 is cytosolic, while STMN2, 3 and 4 share a doubly palmitoylated N-terminal extension that targets them to endomembranes [Gilbert Di Paolo et al., 1997; Lutjens et al., 2000; Poulain and Sobel, 2007]. STMN1 is expressed ubiquitously, STMN2 and STMN3 primarily in neurons, STMN4 in oligodendrocytes, and STMND1 in multiciliated epithelia [Bièche et al., 2003].

Tubulin binding to stathmin SLDs has been measured biochemically and confirmed by atomic resolution structures [Deng et al., 2024; Jourdain et al., 1997; Curmi et al., 1997; Gigant et al., 2000; Honnappa et al., 2003]. The curved conformation of the 1:2:2 complex between SLDs and αβ-tubulins explains how stathmins inhibit microtubule polymerization by sequestering tubulin and promoting catastrophes [Steinmetz et al., 2000]. STMN1, the only member of stathmin family that contains only SLD without the membrane-binding N-terminal extension, is believed to act as a negative regulator of microtubule polymerization by binding to tubulin during mitosis and interphase signaling. Its tubulin-binding activity is regulated by phosphorylation [Brattsand et al., 1994; Marklund et al., 1996; Steinmetz et al., 2001; Niethammer et al., 2004]. Many studies generalized this microtubule-centered cellular functions to all stathmin family members. However, tubulin binding by the membrane-associated stathmins (STMN2, STMN3, STMN4 and STMND1) was measured using only the SLD, with the membrane-targeting N-terminal extension removed [Charbaut et al., 2001; Gigant et al., 2000]. It was unclear whether these stathmins can bind membranes and tubulin simultaneously or whether their membrane-bound form has some function other than destabilizing microtubules. One purpose of this study was to address these question for STMN2.

Two recent studies brought STMN2 into focus in neurological disease research by identifying it as a downstream effector of TDP-43 dysfunction, which is implicated in 97% of ALS cases [Klim et al., 2019; Melamed et al., 2019]. Both studies demonstrated that TDP-43 dysfunction leads to cryptic mis-splicing of the human STMN2 transcript, leading to downregulation of STMN2 protein levels. Subsequent pathology studies revealed that truncated STMN2 transcripts could serve as biomarkers of TDP-43 dysfunction in ALS and frontotemporal dementia [Prudencio et al., 2020]. Collectively, these studies revealed a neuroprotective function of STMN2 which is lost in ALS.

STMN2 was initially identified as an upregulated protein during neuronal development [Grenningloh et al., 2004; Leonard et al., 1987; Pellier−Monnin et al., 2001; Riederer et al., 1997], It is not essential for mice to reach adulthood. However, abnormal compartmentalization and metabolism of STMN2 in the brain and motor neuron-associated tissues are linked to neurodegenerative diseases [Pellier−Monnin et al., 2001; Guerra San Juan et al., 2022; Krus et al., 2022; Mori and Morii, 2002; Himi et al., 1994]. STMN2^-/-^ mice develop late-onset motor neuron degeneration, supporting a neuroprotective role for STMN2 [Guerra San Juan et al., 2022; Krus et al., 2022]. Replenishing STMN2 in affected neurons has been shown to restore neurite outgrowth, prompting the development of clinical trials using splice-correcting oligonucleotides [Baughn et al., 2023]. In-depth understanding of STMN2’s role could help uncover the pathological mechanisms and define therapeutic windows for neurodegenerative diseases. However, the precise molecular function of STMN2 in neurons remains unknown, limiting our understanding of its neuroprotective role. STMN2 is enriched in growth cones and distal stumps following axotomy, suggesting functions in membrane and/or cytoskeleton dynamics [Di Paolo et al., 1997; Shin et al., 2014]. By analogy to STMN1, STMN2 may negatively regulate microtubule. Alternatively, its membrane bound form might regulate vesicle trafficking or serve as a transport adapter, but in that case the purpose of binding tubulin is unclear. Membrane and cytoskeleton functions may combine if STMN2 is a tubulin transport adapter, in which case STMN2 would need to bind tubulin and membranes simultaneously.

A key insight into STMN2 regulation—and potentially its function—is its rapid turnover, with a half-life of less than two hours in cultured neurons [Shin et al., 2012; Thornburg-Suresh et al., 2023]. This characteristic may explain why STMN2 protein level is relatively low compared to its mRNA level in mature neurons [Mori and Morii, 2002]. Notably, the remarkably fast degradation rate of STMN2 [Shin et al., 2012; Thornburg-Suresh et al., 2023], is significantly faster than most of the neuronal proteins [Dörrbaum et al., 2018; Fornasiero et al., 2018], and unusually rapid even for a membrane-associated protein [Dörrbaum et al., 2018]. This raises a key question: how is the homeostasis of STMN2 regulated across different stages of neuronal development despite its rapid turnover?

Here, we analyzed the mechanisms of STMN2 turnover in neurons and a surrogate cell type and used turnover rate to infer its interaction with tubulin in living cells. We found that STMN2’s N-terminal membrane-binding domain is responsible for rapid turnover, while its tubulin-binding SLD stabilizes the non-membrane-bound form of STMN2. Pull-down assays suggested that STMN2 binds either tubulin or membranes, but not both simultaneously. While our study does not reveal the molecular function of STMN2 required to maintain neuronal homeostasis, it suggests this function resides in the membrane-bound, tubulin-free form, with tubulin binding regulating STMN2 stability and location.

## RESULTS AND DISCUSSION

### STMN2 undergoes rapid turnover by the ubiquitin-proteasome pathway in two cellular systems

To determine the turnover rate of STMN2, we performed a cycloheximide (CHX) chase assay in two systems: cancer-derived U2OS cells artificially expressing a V5-tagged STMN2 construct (based on the canonical sequence UniprotID Q93045) (Fig. 1A) under a doxycyclin-inducible Tet-On system, and transcription-factor (TF)-induced pluripotent stem cell (iPSC)-derived neurons (iNs), resembling cortical excitatory neurons [Busskamp et al., 2014]. U2OS cells were more tractable for biochemical assays, while the neuronal system is more *in vivo* relevant. In the U2OS system, STMN2-V5 migrated as two closely-spaced bands on SDS-PAGE, detected by both anti-V5 and anti-STMN2 antibodies. When the inducing reagent was removed, STMN2-V5 levels remained stable for the several hours (Fig. 1B). When inducing reagent was removed and translation was inhibited by CHX, STMN2-V5 bands rapidly decayed with an estimated half-life of approximately 2 hrs (Fig. 1B). STMN2 mRNA was undetectable in iPS cells but was strongly induced by neuron-specifying TFs (see RNA seq data in ref [Busskamp et al., 2014]). We performed turnover assays when endogenous STMN2 expression peaked, a few days after initiating differentiation (ref [Busskamp et al., 2014], Fig. 1C and Supplementary Fig. 1). STMN2 levels were measured via western blot using STMN2-specific antibodies. Endogenous STMN2 showed a similar half-life (∼2 hrs) in iNs (Fig. 1D), similar to previous studies in rodent primary neurons [Shin et al., 2012; Thornburg-Suresh et al., 2023].

**Fig 1.**
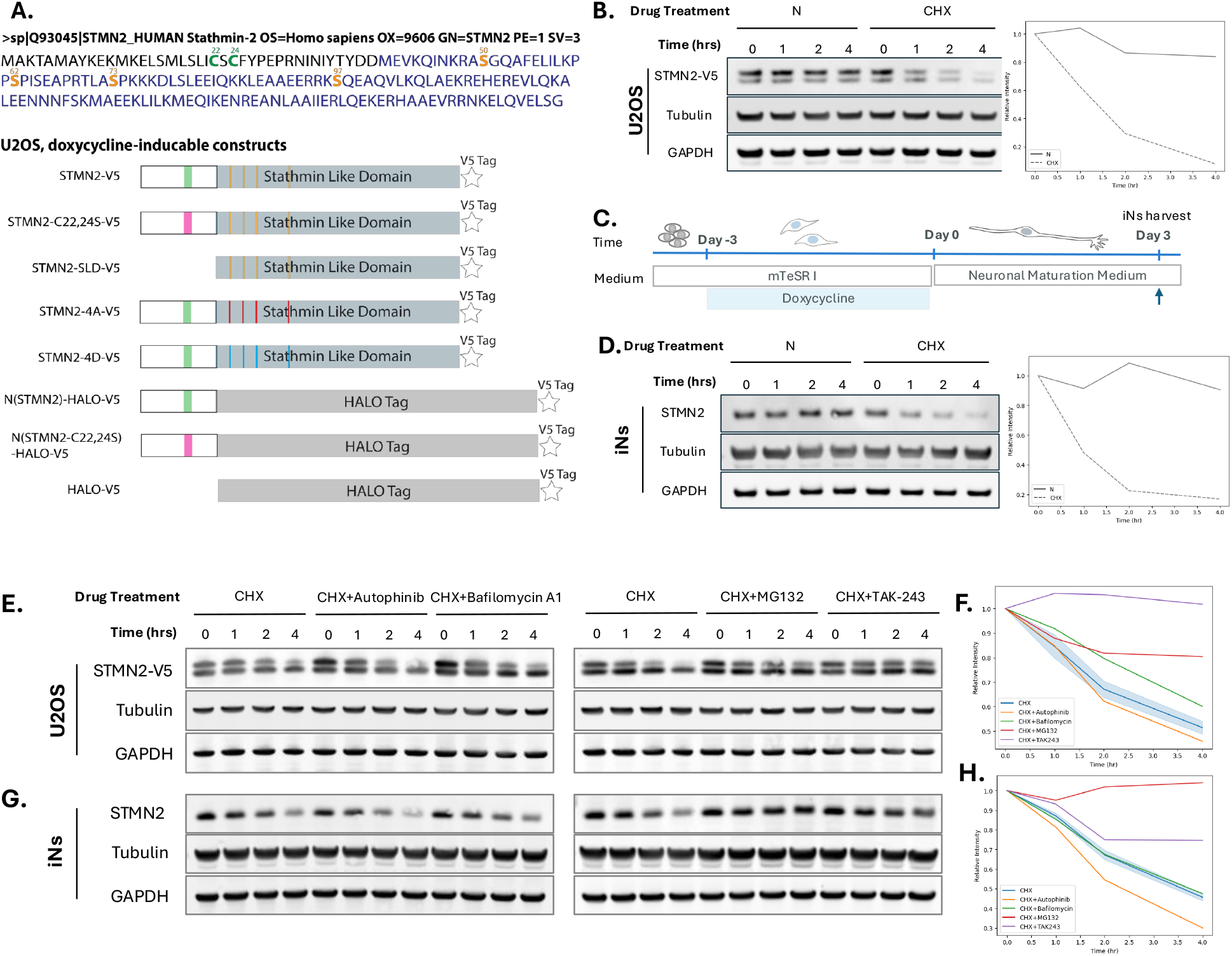
STMN2 undergoes rapid turnover in two cellular systems, with degradation primarily mediated by the ubiquitin-proteasome pathway. **A**. The design of doxycycline-induced constructs with V5 tag. Top, The canonical amino acid sequence of STMN2 (Q93045) used as the basis for the constructs. The Stathmin-like domain (SLD) is highlighted in blue. Two palmitoylatable cysteines (C22, C24) are marked in bold green. Four phosphorylatable serine sites (S50, S62, S73, S97) are marked in bold orange. Bottom, Illustration of the constructs. Palmitoylation sites are shown in green. Magenta indicates the abolished palmitoylation sites, where cysteines are mutated to serines. The four phosphorylatable serine sites (S50, S62, S73, S97) are represented by vertical orange lines. The non-phosphorylatable mutant, where serines are substituted with alanines (4A), is marked in red. Phosphorylation-mimicking mutants, where serines are substitute with aspartate residues, are shown in blue. **B**. Western blot results of the CHX chase assay in U2OS cells with inducible STMN2-V5 expression. STMN2-V5 expression was induced overnight with doxycycline. For the CHX chase assay, the doxycycline-containing media was replaced with either fresh media or CHX-containing media. The start of the assay was defined as the point when the media was changed.. ∼35 μg of whole cell lysate were loaded to each lane. Left panel, STMN2 was detected using an anti-V5 antibody. Right panel, the intensity of STMN2 bands was measured after background subtraction and compared to the starting time point. Left panel, the relative intensity of each band under both conditions was plotted against chase time. **C**. The timeline and experimental system of stem cell derived neurons. The expression of neuronal transcription-factor, NGN1, was induced by doxycycline for three days to generate neurons. A CHX chase assay was conducted on the third day after plating. **D**. Western blot results of CHX chase assay on endogenous STMN2 in transcription-factor (TF)-induced pluripotent stem cell (iPSC) derived neurons (iNs). The start of the assay was defined as the point when CHX was added to the media. A negative control was harvested simultaneously with the CHX-treated neurons for comparison. ∼70 μg of whole cell lysate were loaded to each lane. Left panel, STMN2 was detected using a home-made polyclonal rabbit anti-STMN2 antibody. Right panel, the intensity of STMN2 bands was measured after background subtraction and compared to the starting time point. Left panel, the relative intensity of each band under both conditions was plotted against chase time. **E**. Westen blot of CHX chase assay along with 10 μM Autophinib, 200 nM Bafilomycin A1, 5 μM MG132, or 200 nM TAK-243 in STMN2-V5 expressing U2OS cells. The CHX-only condition samples shown on the left and right panels are from the same experiment and were loaded twice as controls for the western blot. **F**. Relative band intensities of STMN2-V5 (A), plotted against chase time. For each condition, band intensities are normalized to the initial time point (0 hr). Data points for the two CHX-only conditions are plotted with confidence intervals. **G**. Westen blot of CHX chase assay along with 10 μM Autophinib, 200 nM Bafilomycin A1, 5 μM MG132, or 200 nM TAK-243 in iNs. The CHX-only condition samples shown on the left and right panels are from the same experiment and were loaded twice as controls for the western blot. **H**. Relative band intensities of endogenous STMN2 from iNs (C), plotted against chase time. For each condition, band intensities are normalized to the initial time point (0 hr). Data points for the two CHX-only conditions are plotted with confidence intervals.

To determine how STMN2 is degraded, we added inhibitors of autophagosomes, H^+^-ATPases, proteasomes and ubiquitination alongside CHX. The autophagosome inhibitor, Autophinib, had no effect on STMN2 turnover, while the H^+^-ATPases inhibitor, Bafilomycin A1, slightly slowed the turnover rate. In contrast, proteasome and ubiquitin activating enzyme inhibitors (MG132 and TAK-243) largely blocked the turnover of both STMN2-V5 expressed in U2OS (Fig. 1E and F) and endogenous STMN2 in iNs (Fig. 1G and H). We concluded that STMN2 is degraded mostly by the ubiquitin-proteasome system.

### The membrane-binding domain of STMN2 is necessary and sufficient for rapid turnover

STMN2 is targeted to membranes via a short, doubly palmitoylated N-terminal domain [Chauvin et al., 2008; Gilbert Di Paolo et al., 1997; Summers et al., 2018]. This domain is absent in STMN1, which is relatively stable in U2OS cells (data not shown). To investigate the role of this domain in rapid turnover, we engineered U2OS cell lines to inducible express two mutants: STMN2-C22, 24S, with the two palmitoylatable cysteines replaced by serines, and STMN2-SLD, a truncated version lacking the N-terminal domain (Fig. 1A). Upon separating the cellular proteins into soluble and membrane fractions, we found that both constructs localized exclusively to the soluble fraction (Fig. 3A). In the 4-hour CHX chase assay, both constructs displayed a single band on the western blots (Fig. 2B), in contrast to the two bands observed for wild-type STMN2 (Fig 1B). The STMN2-SLD construct exhibited no detectable degradation during the assay (Fig. 2B) and showed higher steady-state levels compared to both wild-type STMN2 and STMN2-22, 24S (Fig. 2B i and ii). Despite its soluble state, STMN2-C22,24S mutant degraded at a similar rate to wild-type STMN2 (Fig. 2B). These findings indicate that the N-terminal domain promotes rapid turnover, but membrane targeting is not required.

**Fig. 2.**
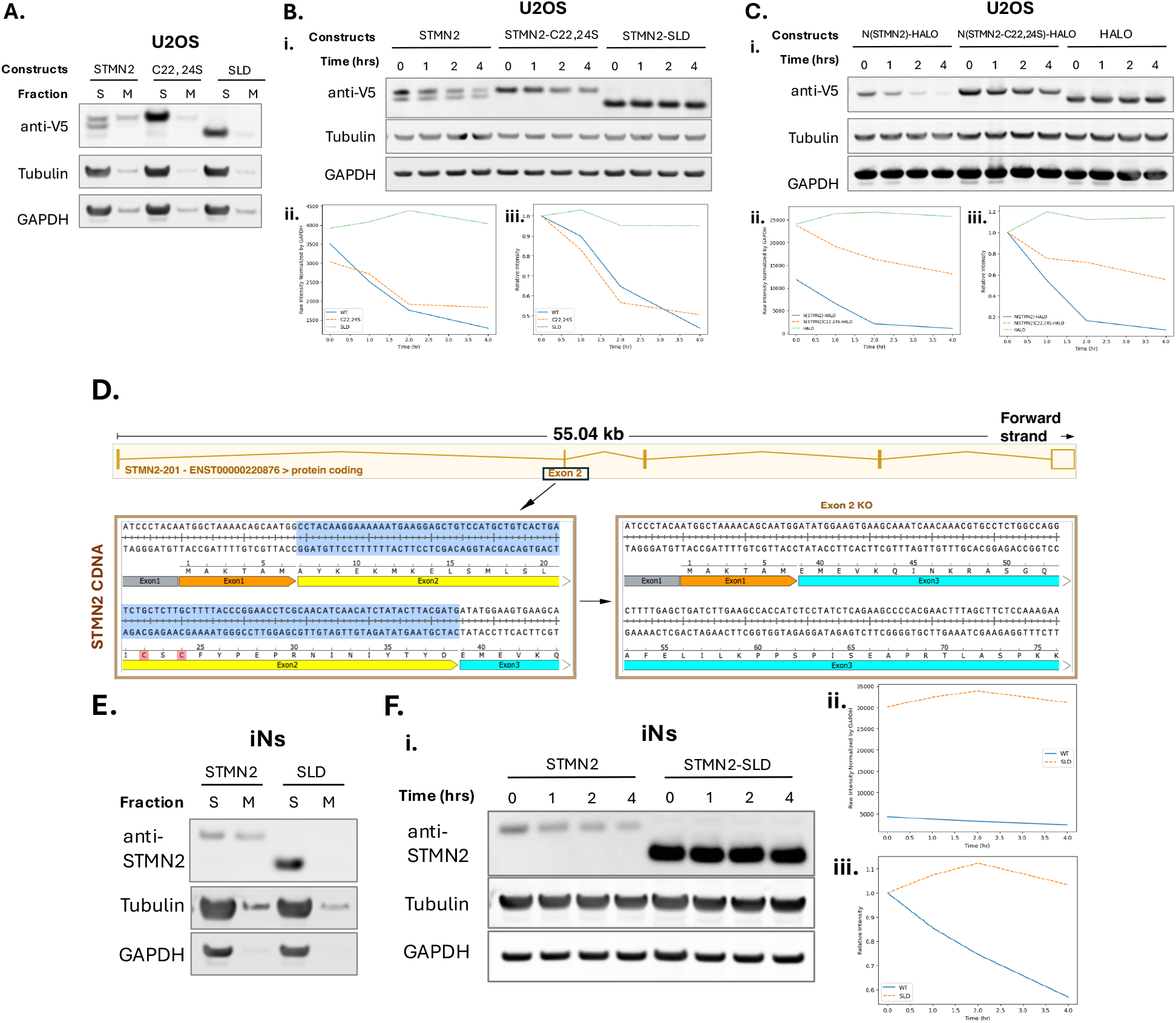
The membrane-binding domain of STMN2 is necessary and sufficient for rapid turnover. **A**. Results of soluble and membrane protein separation experiments from U2OS cell lines expressing wild-type STMN2-V5, STMN2-C22, 24S-V5, and STMN2-SLD-V5. The expression of the proteins was induced overnight with doxycycline. Whole cell lysates were collected by hypotonic shock using low-salt buffer, followed by mechanical disruption by threading through a syringe. Nuclei were removed by low-speed centrifugation. Soluble proteins (S) and membrane proteins (M) were separated by high-speed centrifugation. **B**. CHX chase assay on wild-type STMN2-V5, STMN2-C22, 24S-V5, and STMN2-SLD-V5. Western blot results. **ii**. Raw band intensities of STMN2 constructs normalized by the intensity of corresponding GAPDH are plotted against chase time. **iii**. Relative band intensities plotted against chase time. For each construct, the band intensities are compared to that at the initial time point (0 hr). **C**. CHX chase assay on N(STMN2)-HALO-V5, N(STMN2-C22, 24S)-HALO-V5, and HALO-V5. **i**. Western blot results. **ii**. Raw band intensities of STMN2 constructs normalized by the intensity of corresponding GAPDH are plotted against chase time. **iii**. Relative band intensities plotted against chase time. For each construct, the band intensities are compared to that at the initial time point (0 hr). **D**. CRISPR knock-out scheme for generating a truncated version of STMN2 in iPSCs. Exon 2 of the STMN2 gene was deleted using two sgRNAs that target the flanking regions of Exon 2. The resulting open reading frame lacks Exon 2 (highlighted in yellow), which contains the majority of the N-terminal extension, including the palmitoylatable cysteines responsible for membrane binding. **E**. Results of soluble and membrane protein separation experiments from iNs expressing wild-type STMN2 or truncated STMN2 (SLD). **F**. CHX chase assay on wild-type iNs or iNs expressing STMN2-SLD. i. Western blot results.. **ii**. Raw band intensities of WT-STMN2 or STMN2-SLD detected by anti-STMN2 antibody normalized by the intensity of corresponding GAPDH are plotted against chase time. **iii**. Relative band intensities plotted against chase time. The band intensities of WT-STMN2 or STMN2-SLD are compared to that at the initial time point (0 hr) respectively.

**Fig 3.**
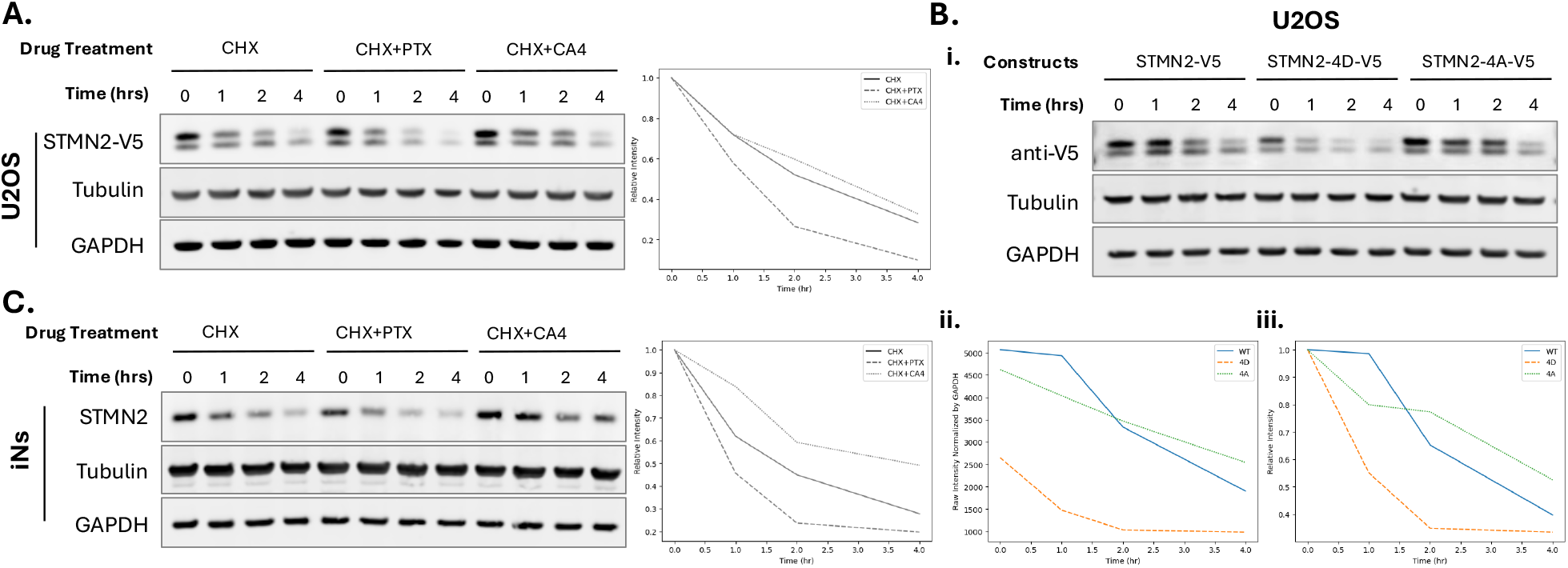
Tubulin binding slows STMN2 turnover. **A**. The degradation of induced STMN2 in the U2OS cell line was influenced by microtubule-stabilizing and depolymerizing drugs. The expression of STMN2-V5 and the CHX chase assay were conducted as described in Figure 1. The microtubule-stabilizing drug paclitaxel (PTX) or the microtubule-depolymerizing drug combretastatin A4 (CA4) was added to the media along with CHX to alter the concentration of soluble tubulin and monitor its effect on STMN2 degradation. **B**. CHX chase assay on wild-type STMN2-V5, phospho-mimetic aspartate mutant (STMN2-4D-V5), and non-phosphorylatable mutant (STMN2-4A-V5). **i**. Western blot results. **ii**. Raw band intensities of STMN2 constructs normalized by the intensity of corresponding GAPDH are plotted against chase time. **iii**. Relative band intensities plotted against chase time. For each construct, the band intensities are compared to that at the initial time point (0 hr). **C**. The degradation of endogenous STMN2 in iNs changed by microtubule polymerization drug or microtubule depolymerization drug. PTX or CA4 was added to the media along with CHX to alter the concentration of soluble tubulin and monitor its effect on STMN2 degradation.

To determine if the N-terminal domain alone promotes rapid turnover, we fused it to HALO tag, a folded domain that does not bind tubulin (N(STMN2)-HALO) (Fig. 1A). We also created a similar fusion using the STMN2-C22, 24S mutant N-terminal domain (N(STMN2-C22, 24S)-HALO) (Fig. 1A). HALO tag alone was stable during CHX treatment (Fig. 2C). In contrast, N(STMN2)-HALO degraded rapidly, with a half-life of approximately 1 hour (Fig. 2C i and iii). The N(STMN2-C22, 24S)-HALO fusion degraded significantly more slowly than N(STMN2)-HALO (Fig. 2C i and iii), starting with a much higher steady-state level (Fig. 2C i and ii). These findings indicate that the N-terminal domain drives STMN2 turnover, with membrane targeting accelerating the process. However, there are additional elements within this short extension (∼39 amino acids) also contribute to STMN2 turnover. The comparable degradation rates of wild-type STMN2 and STMN2-C22, 24S mutant, versus the faster degradation of N(STMN2)-HALO compared to N(STMN2-C22, 24S)-HALO, suggest that the SLD slows turnover compared to HALO, perhaps because it binds tubulin.

To investigate the role of the N-terminal domain of wild-type STMN2 expressed at endogenous levels in iNs, we used CRISPR technique to delete exon 2 of the STMN2 gene in iPSCs, generating a truncated STMN2 variant that retains only the SLD while remaining under the control of endogenous promotor (Fig. 2D). When separating soluble and membrane proteins of iNs expressing either wild-type or truncated STMN2, the truncated STMN2 was found exclusively in the soluble fraction, whereas wild-type STMN2 localized to both soluble and membrane fractions (Fig. 2E). The truncated STMN2 exhibited homeostasis level approximately five times higher than that of the wild-type STMN2 (Fig. 2F i and ii). A 4-hour CHX chase assay revealed no detectable turnover of the truncated STMN2, in stark contrast to the rapid turnover observed with wild-type STMN2 (Fig. 2F). Hence, the difference in homeostasis levels of wild-type and truncated STMN2 is likely attributable to difference in their turnover rates. These findings confirm that the N-terminal domain is critical for promoting the rapid turnover of STMN2, consistent with results from the artificial expression system.

Interestingly, when the panel of inhibitors used in Fig 1E-H was applied to N(STMN2)-HALO-V5 (Fig. 1A and Supplementary Fig. 2A), its turnover, which was significantly faster than that of full-length STMN2, was only slightly slowed down by Bafilomycin, MG132 and TAK-243 (Supplementary Fig. 2). This suggests that the SLD inhibits STMN2 degradation and the N-terminal domain engages additional, non-canonical degradation pathways. In summary, the N-terminal domain of STMN2 appears to promote STMN2 turnover through multiple pathways.

### Tubulin binding slows STMN2 turnover

We next asked if tubulin binding regulates STMN2 turnover. To modulate the level of soluble tubulin within a physiological range, we applied a drug strategy reported previously [Deng et al., 2024]. The microtubule depolymerizing drug (combrestatin A4, CA4) was used to increase soluble tubulin, and the microtubule stabilizing drug (paclitaxel, PTX) to decrease it. The stathmin SLD is known to bind well to tubulin with drugs bound in the colchicine site such as CA4 [Jourdain et al., 1997; Ravelli et al., 2004]. We then measured STMN2 turnover rate in U2OS cells and iNs under microtubule drug treatment. CA4 significantly slowed the turnover of STMN2 in both cell types, whereas PTX accelerated it (Fig. 3A and C). These observations show that tubulin binding to the SLD stabilizes STMN2.

We next modulated tubulin binding by mutating key phosphorylation sites in STMN2. Phosphorylation on conserved serine residues negatively regulates STMN2’s interaction with tubulin and tubulin binding affinity can be experimentally modulated using serine (S) to alanine (A) or aspartate (D) mutations [Antonsson et al., 1998; Chang et al., 2003; Tararuk et al., 2006; Westerlund et al., 2011]. We generated two STMN2 constructs, mutating S50, S62, S73, and S97 to either alanine or aspartate residues (referred to as 4A and 4D mutants respectively, Fig. 1A), The non-phosphorylatable 4A mutant possesses stronger tubulin-binding affinity, while the phospho-mimetic 4D mutant exhibits lower tubulin affinity [Steinmetz et al., 2001]. When expressed in U2OS cells, the 4D mutant was present a significantly lower concentration in ∼5 mg/ml whole cell lysate, with approximately 60% of the concentration of wild-type or 4A (Fig. 3B i and ii). Additionally, the 4D mutant showed a faster turnover rate (Fig. 3B i and iii) compared to both wild-type and the 4A mutant. In contrast, the 4A mutant exhibited expression level similar to wild-type STMN2 but displayed a slower turnover rate. These data support the drug experiment in showing that tubulin binding slows STMN2 turnover.

### Tubulin binding and Trans-Golgi Network (TGN) localization of STMN2 are inversely correlated

We next used a proximity labeling proteomics approach to identify cellular membranes that recruit STMN2, and to test whether membrane targeting is influenced by tubulin binding. We transfected U2OS cells with an inducible STMN2-APEX2 expression cassette and utilized CRISPR technique to knock in an APEX2 tag at the C-terminal of the STMN2 gene in iPSCs. APEX2 oxidizes biotin probes into short-lived reactive radicals that covalently label nearby amino acids, enabling proteomic identification of proteins in proximity to the APEX2-tagged bait protein [Hung et al., 2014; Rhee et al., 2013]. The overall profile of proteins labeled by STMN2-APEX2 in U2OS cells and iNs was similar, although the labeling in iNs was less pronounced, likely due to the lower expression levels of endogenous STMN2. Proximity labeling identified STMN2 as proximal to various proteins, with cell adhesion proteins standing out as highly significant (Fig. 4A and B). Proteasome complexes, early endosomes, and lysosomal membranes were also emerged as key membrane-bound organelles in proximity with STMN2-APEX2 (Fig. 4A and B)

**Fig 4.**
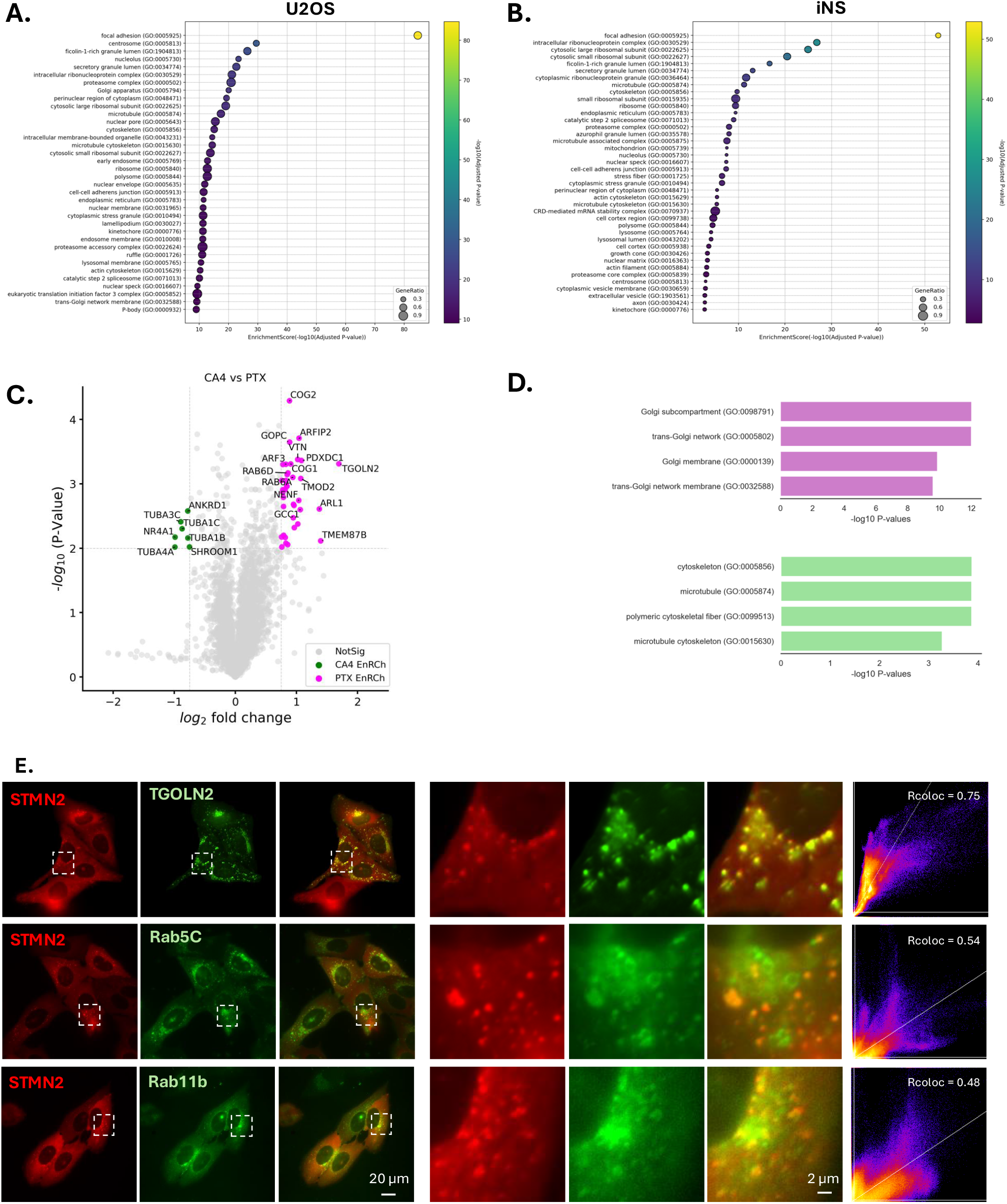
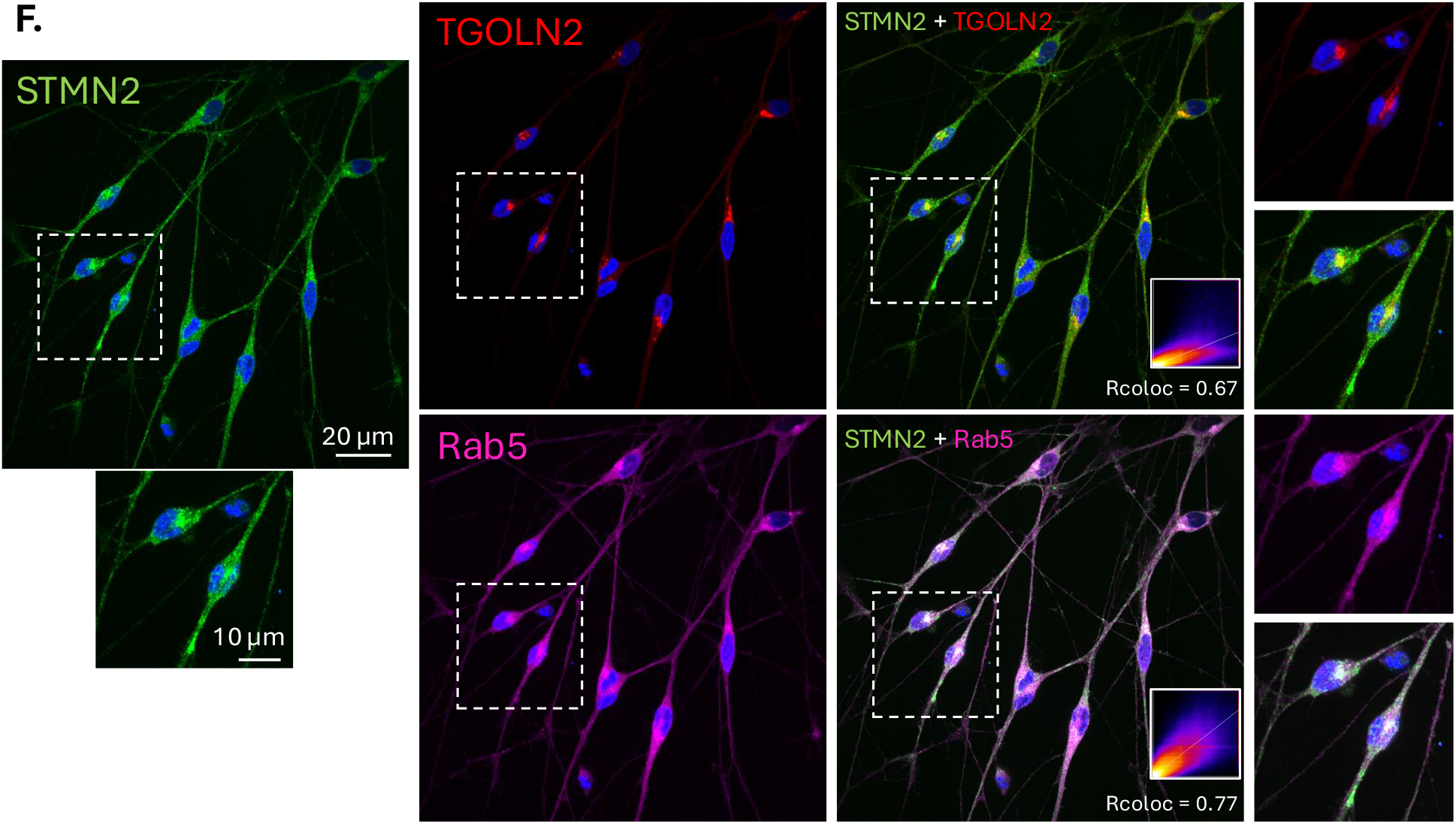
Tubulin binding and Trans-Golgi Network (TGN) association of STMN2 are inversely correlated. **A**. Enrichment gene ontology plot of proximity proteomics in STMN2-APEX2 expressing U2OS. STMN2-APEX2 was induced for 16 hrs prior to the proximity labeling procedure. The size of the dots represents the percentage of genes detected within specific cellular components. The color of the dots indicates the statistical significance of the enrichment. The gene set used is “GO Cellular Component 2018.” **B**. Enrichment gene ontology plot of proximity proteomics of STMN2-APEX2 expressing iNs. D3 iNs was used for this experiment. **C**. Volcano plots showing statistical significance versus protein enrichment in U2OS cells expressing STMN2-APEX2 and treated with CA4 versus PTX. The P values (statistical significance of enrichment calculated from unpaired Student’s t tests) for 3786 quantified proteins were plotted against the TMT ratios of the comparison pair. Magenta and green dots indicate proteins that are increased or decreased in PTX-treated samples, respectively, while grey dots represent proteins with no significant changes. **D**. Gene ontology annotation of the increased and decreased genes upon PTX treatment. (Magenta, PTX increased; green, PTX decreased). **E**. U2OS cells expressing STMN2-mCherry (red) with TGOLN2-EGFP, Rab5C-EGFP, or Rab11b-EGFP (green). STMN2-mCherry expression was induced for 5 hours before live imaging. The areas within the dotted squares are shown in a 10x enlarged view. Colocalization between STMN2 and TGOLN2/Rab5C/Rab11b was estimated using Pearson’s correlation test. The correlation coefficients of pixels above the background are indicated in the plots. **F**. iNs were stained with antibodies against STMN2 (green), TGOLN2 (red), and Rab5 (magenta). The areas within the dotted squares are shown in a 1.5x enlarged view. Colocalization between STMN2 and TGOLN2/Rab5 was estimated using Pearson’s correlation test. The correlation coefficients of pixels above the background are indicated in the plots.

Proximity labeling datasets often identify numerous proteins near a bait. One way to prioritize proteins of interest is to score the effects of molecular perturbation on these interactions. To explore the effect of tubulin binding on proximity labeling, we used the microtubule perturbation strategy discussed above (Fig 4A). PTX, which lowers soluble tubulin, decreased proximity labeling of tubulin and increased the proximity labeling of Golgi apparatus proteins, particularly those in the trans-Golgi network (Fig. 4C and D). CA4, which increases soluble tubulin, had the opposite effect (Fig. 4C and D). The negative correlation between tubulin and trans-Golgi association of STMN2 observed by proximity labeling suggests that these binding events are mutually exclusive. This tubulin-related bidirectional proximity interactomes are reminiscent of observations for STMND1, where tubulin binding inhibited nuclear translocation [Deng et al., 2024].

STMN2 was previously shown to localize to the Golgi apparatus in neurons [Chauvin et al., 2008; Gilbert Di Paolo et al., 1997; Lutjens et al., 2000], consistent with our tubulin-regulated proximity data. To test this localization by imaging, we generated U2OS cell lines expressing STMN2-mScarlet along with GFP-tagged version of three hits from proximity proteomics dataset which are also well-known compartment markers: TGOLN2-GFP (TGN), Rab5C-GFP (late endosomes) and or Rab11b-GFP (recycling endosomes). Live-cell imaging revealed that STMN2 exhibited the strongest colocalization with TGOLN2 (Pearson’s correlation coefficient, Rcoloc = 0.75), with partial colocalization observed with Rab5C (Rcoloc = 0.54) and Rab11b (Rcoloc = 0.48) (Fig. 4E.).

To perform a similar co-localization analysis in neurons, we stained iNs with antibodies against STMN2, TGOLN2 and Rab5A. TGOLN2 was primarily confined to cell bodies, while Rab5A scattered throughput iNs. In neuronal cell bodies, STMN2 strongly colocalized with TGOLN2 (Rcoloc = 0.67) (Fig. 4F.). However, overall co-localization was stronger with Rab5A, driven by the presence of Rab5A in axons where it partially colocalized with STMN2 (Rcoloc = 0.77) (Fig. 4F.). These findings confirm that STMN2 localizes to the TGN in the cell body of neurons but suggest a more complex localization in axons.

### Tubulin binds more strongly to soluble than membrane-bound STMN2

A key conceptual question for STMN2 regulation and function is whether tubulin binding and membrane binding can occur simultaneously or are mutually exclusive as suggested by our proximity proteomics. To address this, we firstly used imaging to examine whether tubulin and STMN2 co-localize on membranes in transfected U2OS cells or iNs. We assessed both endogenous and tagged proteins, using various fixation or live imaging methods. Significant co-localization between tubulin and STMN2 was not observed under any condition. An example of live-imaging in iNs is shown in Supplementary Fig. 3. Negative results from co-localization assays could have multiple interpretations, but they align with the hypothesis that tubulin binding and membrane targeting of STMN2 are mutually exclusive.

To directly compare tubulin binding by soluble vs membrane-bound STMN2 we developed a pull-down strategy. U2OS cells with or without STMN2-V5 expression were subjected to hypotonic lysis without detergent and separated into cytosol and membrane fractions by centrifugation. To preserve weak protein interactions, some samples were treated with zero-length cross-linking using 1-ethyl-3-(3-dimethylaminopropyl)carbodiimide (EDC) before immunoprecipitation. This reagent was previously shown to efficiently cross link the SLD of STMN1 to tubulin [Curmi et al., 1997; Steinmetz et al., 2000]. Detergent was then added to both fractions, and STMN2 was immunoprecipitated utilizing the V5 tag. Immunoblots were probed with antibodies against V5 and tubulin. Inspection of gels revealed substantially more tubulin co-precipitating with STMN2 from the cytosol fraction compared to the membrane fraction, but more STMN2 was also recovered from this fraction. To normalize the degree of co-precipitation we quantified the intensity of bands, subtracted local background, and accounted the minor tubulin signal observed in the immunoprecipitation from control samples without STMN2. Based on this analysis, soluble STMN2 bound approximately 10-fold more tubulin per unit of STMN2 than membrane-bound STMN2 (Fig. 5A and B). This result was unaffected by cross-linking. Cross-linking generated a 70 kDa band, consistent with the previously reported STMN1-tubulin cross-link [Curmi et al., 1997; Steinmetz et al., 2000]. Higher molecular weight bands in the EDC-treated samples presumably correspond to varying degrees of cross-linking of the 1:2:2 complex between stathmins and tubulin. In contrast, minimal tubulin cross linking was observed in the membrane fraction (Fig. 5A and B). Taken together with the proximity labeling and lack of co-localization, these findings support the hypothesis that tubulin and membrane binding by STMN2 are mutually exclusive.

**Fig 5.**
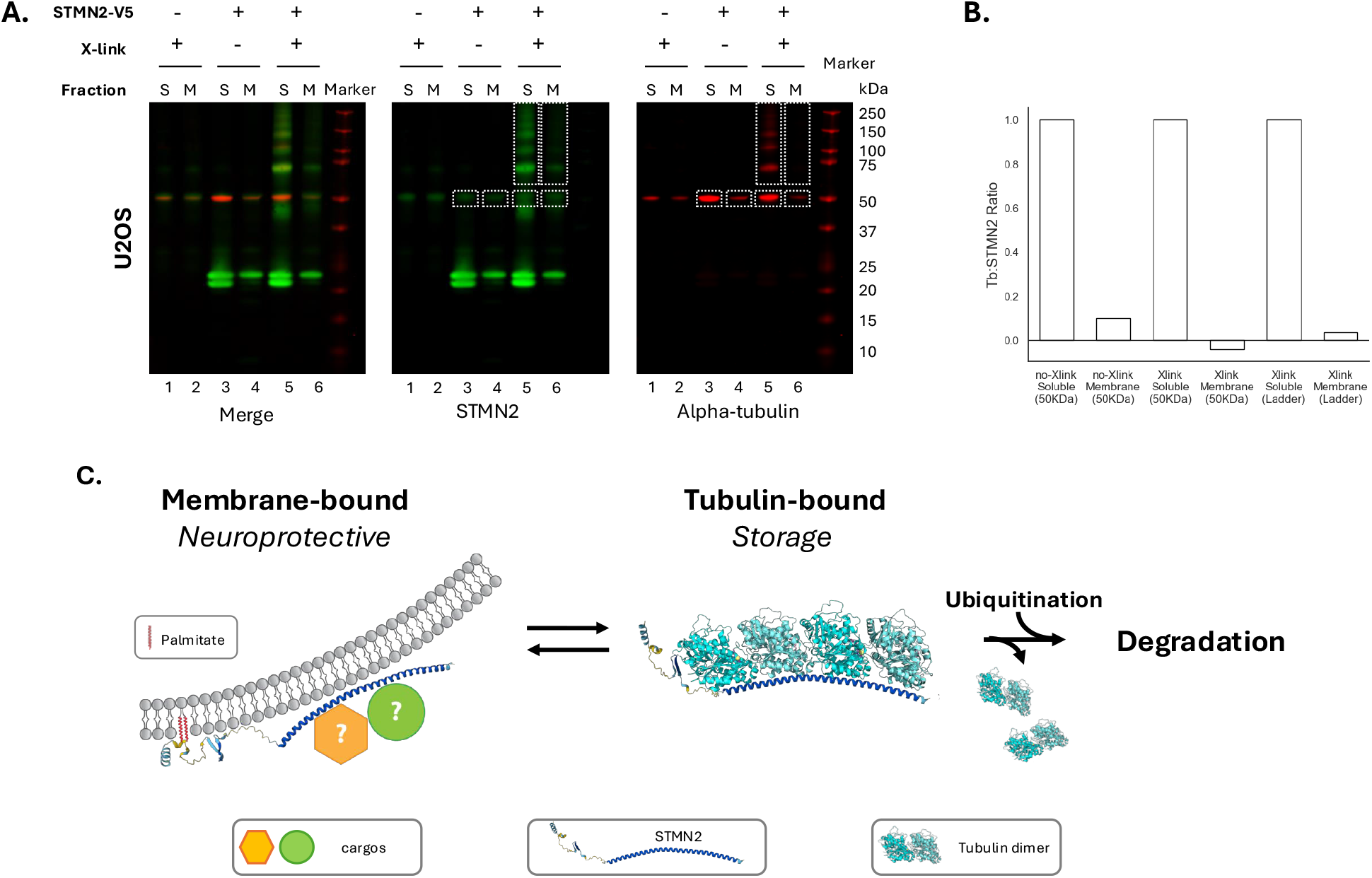
Tubulin prefers binding to soluble than membrane-bound STMN2. **A**. Western blot of Immunoprecipitation of U2OS cells with or without STMN2-V5 expression with antibody against V5 tag. The membrane is blot using antibody against STMN2 (red) and antibody against alpha-tubulin (green). The cell lysate was prepared as described in Figure 2. From left to right, U2OS cells without STMN2-V5 expression, cross-linked before immunoprecipitation (Lane 1, soluble fraction, S; lane 2, membrane fraction, M). U2OS cells expressing STMN2-V5, no cross-linking before immunoprecipitation (Lane 3, soluble fraction, S; lane 4, membrane fraction, M). U2OS cells expressing STMN2-V5, cross-linked before immunoprecipitation (Lane 5, soluble fraction, S; lane 6, membrane fraction, M). **B**. Plot of intensity ratio of tubulin signal vs STMN2 signal. The measured areas are marked by white dotted blocks. Background is subtracted before calculating ratio. From left to right: 50 KDa band of soluble fraction of sample without cross-linking step (lane 3); 50 KDa band of membrane fraction of sample without cross-linking step (lane 4); 50 KDa band of soluble fraction of crosslinked sample (lane 5); 50 KDa band of membrane fraction of crosslinked sample (lane 6); ladder area of soluble fraction of crosslinked sample (lane 5); ladder area of membrane fraction of crosslinked sample (lane 6). **C**. The proposed model of STMN2’s fast turnover. STMN2 interconverts between a tubulin-free, membrane-bound form and a tubulin-bound, soluble form. The binding partners of membrane-bound STMN2 is unknown. When the soluble form loses bound tubulin, STMN2 becomes rapidly ubiquitinated and degraded. STMN2’s poorly understood neuroprotective function possibly depends on some unknown molecular activity of the membrane bound form. The structure of STMN2 is based on AF-Q93045-F1-V4 from Alpha Fold. The state of palmitoylation of tubulin-bound STMN2 is not considered.

## Discussion

The most pressing question in stathmin research is to determine the molecular function of STMN2 that protects motor neurons from degeneration and, more generally, to understand the molecular function of membrane-targeted stathmins. Here, we sought to better characterize STMN2 and in particular to determine the function of its N-terminal membrane targeting domain. The best-characterized molecular activity of STMN2 is its ability to bind tubulin and inhibit microtubule polymerization in biochemical assays. However, these activities were only demonstrated for expressed SLDs lacking the N terminal membrane binding domain. Whether full length STMN2 binds tubulin, and whether negative regulation of MTs is the primary cellular function of STMN2, were unclear. An alternative possibility, which we currently favor, is that tubulin binding regulates STMN2, while its neuroprotective function depends on some as-yet-unknown activity of its membrane-bound form.

As a start towards distinguishing these hypothesis, we used STMN2 turnover rate as a proxy for its molecular interactions in living cells. We then applied two independent approaches to perturb membrane and tubulin binding, STMN2 mutations and drug treatments that alter the concentration of soluble tubulin. We found that tubulin binding stabilizes STMN2 against degradation, while rapid degradation is primarily caused by the N-terminal domain (Fig. 1-3). These results suggest that, rather than STMN2 regulating tubulin and microtubule as usually proposed, tubulin binding may instead serve as a regulatory mechanism for STMN2 function.

An important clue to STMN2 function is its membrane localization. When membrane-bound, may regulate the dynamics of its target membrane, or serve as a transport adapter of membrane-bound cargos. Previous studies reported that STMN2 localized primarily to the Golgi in both neurons and transfected cells [Chauvin et al., 2008; Gilbert Di Paolo et al., 1997; Lutjens et al., 2000], but the identity of the membranes which recruit STMN2 in axons and synapses is unclear, with some reports suggesting an association with mitochondria under certain conditions [Chauvin et al., 2008]. To refine our understanding of STMN2 localization in both U2OS cells and iNs, we used STMN2-APEX2 proximity labeling (Fig. 4A and B) and validated interactors through live-cell and immunofluorescent imaging (Fig. 4E and F). Our data confirm that STMN2 preferentially localizes to the TGN, but also reveal complexities in its neuronal localization. Although our proximity labeling approach failed to identify specific interaction partners, possibly due to STMN2’s proximity to too many proteins, comparing interactomes with and without microtubule-targeting drugs established a clear negative correlation between interacting with tubulin and TGN membranes. This finding further supports our hypothesis that membrane-binding and tubulin-binding are mutually exclusive.

To directly test if tubulin and membrane interactions of STMN2 are mutually exclusive, we performed IP-western experiments on cell lysate, with or without a cross linker to stabilize the interaction with tubulin (Fig. 5A and B). The results of these experiments showed that tubulin in cell lysate preferentially associates with the soluble form of STMN2. While a small amount of tubulin was detected in immunoprecipitants from the membrane fraction, the sensitivity of the assay was limited by non-specific IP of tubulin measured in cells lacking STMN2 expression. This raises the possibility that tubulin does not actually bind to membrane-associated STMN2. However, to confirm this, further validation using pure protein assays is required. This future direction will involve the purification of doubly palmitoylated STMN2 and its reconstitution into synthetic membranes, which has not been reported and likely to prove technically challenging.

Our results and their implications are summarized in Fig. 5C. Our data suggest that STMN2 binds to either membranes or tubulin, but not to both at the same time. In its soluble form, STMN2 is protected from degradation by its association with tubulin. When tubulin dissociates, STMN2 undergoes rapid ubiquitination and degradation. In Fig. 5C, we hypothesize that when tubulin-free STMN2 resides on membranes, its SLD recruits some unknown factor(s), and that this interaction is central to its neuroprotective function. While these unknown proteins implicated in this cartoon are likely present in our proximity labeling datasets, we currently lack the necessary information to distinguish them from less important interactors. In the absence of this knowledge, we can only speculate on STMN2’s neuroprotective function. One possibility is that STMN2 could regulate the dynamics of the vesicles to which it is recruited, e.g. target them towards sites of plasma membrane damage in the axon or synapse. Alternatively, STMN2 could serve as a transport adapter for some factors other than tubulin, which are crucial for neuroprotection, such as proteins or mRNAs needed for membrane repair in the axon or synapse. In both models, tubulin binding serves as an upstream regulator that negatively influences STMN2’s membrane targeting. The concentration of soluble tubulin likely varies across different axonal regions, e.g. it may be lower in axons compared to cell bodies and synapses. This variation suggests that soluble tubulin could serve as a location-dependent regulator of STMN2, influencing the dissociation of its unknown cargos at synapses or sites of axonal plasma membrane damage.

## MATERIAL AND METHODS

### Molecular biology and construct of cell lines

All STMN2 constructs are based on canonical human STMN2 sequence (UniprotID Q93045). All primers and DNA blocks were ordered from Integrated DNA Technologies. The doxycycline-inducible plasmids were based upon a piggyBac vector, PB-TRE-EGFP-EF1a-rtTA (Addgene #104454), and synthesized DNA cassettes were cloned between NcoI and XhoI. Each construct was transfected together with a transposase vector at ratio of 4:1, using Lipofectamine 3000 (Thermo Fisher Scientific, L3000008). Transfected cells were further selected with 1 μg/ml puromycin (Thermo Fisher Scientific, A1113803).

We generated STMN2 exon2 knock-out cell lines using CRISPR [Hsu et al., 2013] system. The sgRNA targeting the exon2 and its flanking regions were designed according to CRISPick [Doench et al., 2016] and CHOPCHOP [Labun et al., 2019]. We inserted a pair of sgRNAs into two expression vectors for sgRNAs and for expression of Cas9 linked to mCherry (Addgene #64324) or GFP (Addgene #48138). The two vectors were transfected into cells simultaneously using Lipofectamine 3000. Transfected cell lines were initially selected by flow cytometry on the expression of both mCherry and GFP, followed by screening of purified genomic DNA (Invitrogen, K182001) using PCR, and finally the genomic loci including the targeted fragments were sequenced.

### Antibodies

Antibodies against the following antigens were used: STMN2 polyclonal rabbit antibody (Invetrogen, PA5-23049); STMN2 monoclonal mouse antibody (Proteintech, 67204-1-Ig); V5 epitope tag rabbit antibody (Novus Biologicals, NB600-381); V5 epitope tag mouse antibody (E10/V4RR) (Novus Biologicals, NBP2-37825); anti-α-tubulin antibody (DM1A) (Sigma-Aldrich, 05-829); GAPDH (D16H11) XP rabbit mAb (Cell Signaling Technology, #5174); Anti-TGOLN2 antibody produced in rabbit (Sigma Aldrich, HPA012723); Rab5A (E6N8S) Mouse mAb (Cell Signaling Technology, #46449); Rab11 (D4F5) XP^®^ Rabbit mAb (Cell Signaling Technology, #5589).

To generate anti-STMN2 serum, rabbits were immunized with 1 mg of synthetic N-terminal-acetylated peptides (GenScript) corresponding to the C-terminal sequence of STMN2 (CRHAAEVRRNKELQVELSG). Boosts were administered every three weeks with 50% the initial amount of peptide in Freund’s adjuvant. Specific antibodies were purified from the anti-STMN2 serum using a STMN2 affinity column, following the manufacturer’s instructions (Pharmacia Biotech). The purified antibody was validated by staining STMN2-V5 expressed in cells with a polyclonal STMN2 rabbit antibody (Invitrogen, PA5-23049), a monoclonal STMN2 mouse antibody (Proteintech, 67204-1-Ig), a V5 epitope tag rabbit antibody (Novus Biologicals, NB600-381), and a V5 epitope tag mouse antibody (E10/V4RR, Novus Biologicals, NBP2-37825). Validation also included co-staining endogenous STMN2 expressed in neurons using the STMN2 monoclonal mouse antibody (Proteintech, 67204-1-Ig).

### Cell culture and inducible expression of STMN2 constructs in U2OS cells

U2OS cells were cultured in DMEM (Corning, 10-013-CV) supplemented with 10% FBS (Gibco, 26140-079). The STMN2 transgene cells were maintained with a final concentration of 1 μg/ml puromycin (Thermo Fisher Scientific, A1113803). The expression of STMN2 constructs in U2OS cells was induced by treatment with 2 μg/ml doxycycline (Sigma, D3072) for 4 hrs for live-imaging or 16 hrs for biochemical analysis.

### Stem cell culture and iNs generation

hiPSC cell lines were generated from WTC-11 cell line (Coriell Institute, GM25256). Neuro-progenitor cells were derived from WTC-11 by transfecting the cells with a piggy-BAC-neurogenin1(NGN1) system kindly provided by George Church lab [Busskamp et al., 2014]. Stem cells were cultured on Matrigel-coated surfaces (Corning, 354277), using mTeSR1 media (STEMCELL™ Technologies, 05850). The NGN1-transformed hiPSC cells lines were selected and maintained in mTeSR1 supplemented with 1 μg/ml puromycin (Thermo Fisher Scientific, A1113803). For iPSC culture, media was changed everyday. For passaging, cells were dissociated using Accutase (STEMCELL™ Technologies, 07920), and replated in mTeSR1 supplemented with 1 μg/ml Y-27632 Rho Kinase inhibitor (Cayman Chemical, 10005583). To induce NGN1 expression, doxycycline (Sigma, D3072) was added at a concentration of 2 μg/ml. After three days of induction, the iNs were dissociated and replated onto poly-d-lysine (Sigma, P6407) and laminin (Thermo Fisher, 23017015) coated surface, using neurobasal A media (Gibco, 10888022) supplemented with B-27 (Gibco, 17504044), MEM Non-Essential Amino Acid (Gibco, 11140050) and GlutaMAX (Gibco, 35050061). For iNS culture, media was changed every 2 days.

### CHX chase assay and immunoblotting

After the expression of STMN2 constructs were induced for 16 hrs in U2OS cells, the induction media was replaced with fresh media containing 20 μg/ml cycloheximide (CHX), and the cells were incubated for 1, 2, or 4 hrs. To collect the cell lysates from CHX chase assay, the cells were washed with cold PBS, harvested, and then lysed using cold EDTA-free NP-40 buffer (50 mM Tris-Cl, pH 8.0, 150 mM NaCl, 1% NP-40 substitute, EDTA-free complete protease inhibitor (Roche, 11836170001)), while rocking for 15 minutes at 4°C. Each well of cell culture in a 6-well plate was lysed with 90ul of NP-40 buffer, achieving a final protein concentration of ∼5 mg/ml. Approximately 35 μg of protein was loaded onto each lane of an SDS-PAGE gel for optimal detection of all STMN2 constructs. After electrophoresis, the gels were transferred onto 0.2 μM nitrocellulose membrane (Bio-Rad) and probed with the following antibodies: V5 epitope tag rabbit antibody (Novus Biologicals, NB600-381); V5 epitope tag mouse antibody (E10/V4RR) (Novus Biologicals, NBP2-37825); anti-α-tubulin antibody (DM1A) (Sigma-Aldrich, 05-829); GAPDH (D16H11) XP rabbit mAb (Cell Signaling Technology, #5174).

For iNs, after three days of plating, CHX was added to the media, and the iNs were incubated for 1, 2, or 4 hrs. Cell lysates were collected in the same manner as for U2OS cells. Approximately 70 μg of protein was loaded onto each lane of an SDS-PAGE gel for effective detection of endogenous STMN2. The following antibodies were used to probe the transferred membranes: homemade STMN2 antibody, anti-α-tubulin antibody (DM1A) (Sigma-Aldrich, 05-829), and GAPDH (D16H11) XP rabbit mAb (Cell Signaling Technology, #5174).

### Cytosolic and membrane protein separation

Cells were washed twice with PBS, scraped into PBS, and spun down. The resulting cell pellets were collected, weighed, and resuspended in 2 μL of 0.5X BrB80 buffer (40 mM K-PIPES, 0.5 mM MgCl_2_, 0.5 mM EGTA, pH 6.8) per milligram of cell mass. The suspension was placed on ice for 30 minutes to induce osmotic shock. Following this, the cell lysate was passed through a 25G needle three times and centrifuged at 500 xg for 5 minutes at 4°C to pellet the nuclei. The supernatant was collected and centrifuged again at 1400 xg for 20 minutes at 4°C to separate the soluble and membrane fractions. The pellet was washed once with 0.5X BrB80 buffer and resuspended in 1 μL of 0.5X BrB80 buffer per milligram of cell mass. Protein concentrations in the soluble and membrane fractions were measured using the BCA assay.

### Cross-linking and co-IP

For U2OS samples, ∼350 μg of protein was used per reaction, while 2 mg of protein was used for iNs. To cross-link interacting proteins, 2 mM of the cross-linker 1-ethyl-3-(dimethylamino)propylcarbodiimide (EDC) was added to each reaction and incubated at 4°C for 30 minutes. The reaction was then quenched by adding 50 mM sodium acetate and 50 mM glycine, and incubating for 30 minutes at 4°C.

For each immunoprecipitation reaction, 8 μL of Dynabeads Protein G (Invitrogen, 10004D) was coated with 6 μg of rabbit anti-V5 IgG (Novus Biologicals, NB600-381), according to the manufacturer’s protocol. Protein samples were adjusted to a final concentration of 100 mM KCl with 0.1% Triton X-100 before being added to the coated Dynabeads for immunoprecipitation. Each reaction was incubated with 8 μL of anti-V5 IgG-coated Dynabeads for 1 hour at 4°C. The Dynabeads were washed five times with PBS + 0.1% Triton X-100, and the bound proteins were eluted into 40 μL of SDS-PAGE loading buffer. For western blot, 10 μL of the eluted sample was loaded onto each lane for detection.

### Proximity proteomics

Proximity labeling was performed on cells expressing APEX2-fused STMND1 constructs. After 4 hr of doxycycline-induced expression, biotinyl tyramide (Toronto Research Chemicals, B397770) was added to the media at a final concentration of 0.5 μM, and cells were incubated in the labeling media for 1 hr. To initiate labeling, H_2_O_2_ (Sigma-Aldrich, H1009) was added to a final concentration of 1 mM. Exactly 1 min after H_2_O_2_ treatment, the labeling solution was decanted, and cells were washed three times with cold quenching solution containing 10 mM sodium ascorbate (VWR 95035-692), 5 mM Trolox (Sigma-Aldrich, 238813), and 10 mM sodium azide (Sigma-Aldrich, S2002). The cell lysate was harvested by scraping and cleared by centrifugation at 16,000 × g for 20 minutes, and the resulting pellet was flash-frozen and stored at −80°C until streptavidin pull-down. The cell pellet was lysed using a cell lysis solution (8 M Urea, 100 mM sodium phosphate pH 8, 1% SDS (w/v), 100 mM NH4HCO3, 10 mM TCEP, sterile-filtered). Protein was extracted by adding 55% TCA (Sigma-Aldrich, 91228) at a 1:1 ratio to the lysate and precipitated by centrifugation. The protein pellet was then washed three times with −20°C cold acetone. Protein was subjected to cysteine alkylation using 20 mM iodoacetamide (Sigma-Aldrich, I6125), quenched by 50 mM DTT (Sigma-Aldrich, 43815), and pulled down using streptavidin magnetic beads (VWR, PI88817). A detailed procedure for the protein processing procedure can be found in *Kalocsay, 2019*[Kalocsay, 2019].

Subsequent protein processing procedures and MS analysis were carried out as described. The digested peptides were labeled with TMTpro 16-plex (Thermo Fisher Scientific, A44520) for 1 h. Data collection followed a MultiNotch MS3 TMT method using an Orbitrap Lumos mass spectrometer coupled to a Proxeon EASY-nLC 1200 liquid chromatography system (both Thermo Fisher Scientific). Peptides were searched against a size-sorted forward and reverse database of the *Homo sapiens* reference proteome (Uniprot 03/2021) using SEQUEST (v.28, rev. 12)-based software. Spectra were first converted to mzXML. For the searches, a mass tolerance of 20 p.p.m. for precursors and a fragment ion tolerance of 0.9 Da were used. The search allowed for a maximum of two missed cleavages per peptide.

Carboxyamidomethylation on cysteine was set as a static modification (+57.0214 Da), and oxidized methionine residues (+15.9949 Da) were searched for dynamically. A target decoy database strategy was applied, and a false discovery rate (FDR) of 1% was set for peptide-spectrum matches after filtering by linear discriminant analysis. The FDR for final collapsed proteins was 1%. MS1 data were calibrated post-search, and searches performed again. Quantitative information on peptides was derived from MS3 scans. Quantitative tables were generated requiring an MS2 isolation specificity of >70% for each peptide and a sum of TMT (tandem mass tags) signal:noise ratio (s:n) of >200 over all channels for any given peptide, and then exported to Excel and further processed therein. Proteomics raw data and search results were deposited in the PRIDE archive. The relative summed TMT s:n for proteins between two experimental conditions was calculated from the sum of TMT s:n for all peptides of a given protein quantified.

### Immunofluorescence and imaging

Cells plated on #1.5 coverslips were washed with PBS and crosslinked with freshly made 4% paraformaldehyde and 0.2% glutaraldehyde in PBS for 15 min at room temperature, and washed three times with PBS afterwards. Permeabilization was carried out with 0.1% Triton-X in PBS, 5 min at RT. Cells were blocked with 5% BSA (constituted from powder BSA, Roche Fraction V, sold by Sigma Catalog Number: 10735078001) in PBS for 1 hr at RT. Primary antibodies were diluted in PBS, and cells were incubated with diluted primary antibodies for ∼16 hr at 4°C in a humidified chamber. Cells were then washed three times with in PBS and incubated with fluorescently labelled secondary antibodies, diluted 1:500 in PBS for 1 hr at RT, and washed three times with PBS. The coverslips were mounted on glass slides using VECTASHIELD plus antifade mounting medium with DAPI (Vector Laboratories, H-2000) and left in a dark chamber overnight before imaging.

Images were acquired with a Nikon Ti motorized inverted microscope equipped with Yokogawa CSU-W1 spinning disk confocal at Harvard Nikon Imaging Center, using a 63x/1.40 NA Oil objective lens. The images were processed in ImageJ FUJI for presentation. Images of cilia were then analyzed in a Jupyter Lab environment using pandas, SciPy, NumPy and plotted with matplotlib and seaborn. The pipelines and Jupyter notebooks are available on GitHub.

## ACKNOWLEDGEMENT

The authors thank Dr. Michel Nofal for valuable discussions and generously providing antibodies. We also acknowledge the Nikon Imaging Center at Harvard Medical School (NIC) for providing technical support of imaging. This study was supported by the National Institute of General Medical Sciences, grant R35GM131753 to Timothy Mitchison and by the Cancer Prevention and Research Institute of Texas (grant RR220032), awarded to Marian Kalocsay, who is a CPRIT Scholar in Cancer Research. The authors declare no competing interests.

## Notes

### Competing Interest Statement

The authors have declared no competing interest.

## REFERENCES

Antonsson B, Kassel DB, Paolo GD, Lutjens R, Riederer BM, Grenningloh G. 1998. Identification of in Vitro Phosphorylation Sites in the Growth Cone Protein SCG10 EFFECT OF PHOSPHORYLATION SITE MUTANTS ON MICROTUBULE-DESTABILIZING ACTIVITY. J Biol Chem 273(14):8439–8446.

Baughn MW, Melamed Z, López-Erauskin J, Beccari MS, Ling K, Zuberi A, Presa M, Gonzalo-Gil E, Maimon R, Vazquez-Sanchez S, Chaturvedi S, Bravo-Hernández M, Taupin V, Moore S, Artates JW, Acks E, Ndayambaje IS, Agra de Almeida Quadros AR, Jafar-nejad P, Rigo F, Bennett CF, Lutz C, Lagier-Tourenne C, Cleveland DW. 2023. Mechanism of STMN2 cryptic splice-polyadenylation and its correction for TDP-43 proteinopathies. Science 379(6637):1140–1149.

Bièche I, Maucuer A, Laurendeau I, Lachkar S, Spano AJ, Frankfurter A, Lévy P, Manceau V, Sobel A, Vidaud M, Curmi PA. 2003. Expression of stathmin family genes in human tissues: non-neural-restricted expression for SCLIP. Genomics 81(4):400–410.

Brattsand G, Marklund U, Nylander K, Roos G, Gullberg M. 1994. Cell-cycle-regulated phosphorylation of oncoprotein 18 on Ser16, Ser25 and Ser38. European Journal of Biochemistry 220(2):359–368.

Busskamp V, Lewis NE, Guye P, Ng AH, Shipman SL, Byrne SM, Sanjana NE, Murn J, Li Y, Li S, Stadler M, Weiss R, Church GM. 2014. Rapid neurogenesis through transcriptional activation in human stem cells. Mol Syst Biol 10(11):760.

Chang L, Jones Y, Ellisman MH, Goldstein LSB, Karin M. 2003. JNK1 Is Required for Maintenance of Neuronal Microtubules and Controls Phosphorylation of Microtubule-Associated Proteins. Developmental Cell 4(4):521–533.

Charbaut E, Curmi PA, Ozon S, Lachkar S, Redeker V, Sobel A. 2001. Stathmin Family Proteins Display Specific Molecular and Tubulin Binding Properties*. Journal of Biological Chemistry 276(19):16146–16154.

Chauvin S, Poulain FE, Ozon S, Sobel A. 2008. Palmitoylation of stathmin family proteins domain A controls Golgi versus mitochondrial subcellular targeting. Biology of the Cell 100(10):577–591.

Curmi PA, Andersen SS, Lachkar S, Gavet O, Karsenti E, Knossow M, Sobel A. 1997. The stathmin/tubulin interaction in vitro. J Biol Chem 272(40):25029–25036.

Deng X, Seguinot BO, Bradshaw G, Lee JS, Coy S, Kalocsay M, Santagata S, Mitchison T. 2024. STMND1 is a phylogenetically ancient stathmin which localizes to motile cilia and exhibits nuclear translocation that is inhibited when soluble tubulin concentration increases. MBoC 35(6):ar82.

Di Paolo G, Lutjens R, Osen-Sand A, Sobel A, Catsicas S, Grenningloh G. 1997. Differential distribution of stathmin and SCG10 in developing neurons in culture. Journal of Neuroscience Research 50(6):1000–1009.

Doench JG, Fusi N, Sullender M, Hegde M, Vaimberg EW, Donovan KF, Smith I, Tothova Z, Wilen C, Orchard R, Virgin HW, Listgarten J, Root DE. 2016. Optimized sgRNA design to maximize activity and minimize on-target effects of CRISPR-Cas9. Nat Biotechnol 34(2):184–191.

Dörrbaum AR, Kochen L, Langer JD, Schuman EM. 2018. Local and global influences on protein turnover in neurons and glia. eLife 7:e34202.

Fornasiero EF, Mandad S, Wildhagen H, Alevra M, Rammner B, Keihani S, Opazo F, Urban I, Ischebeck T, Sakib MS, Fard MK, Kirli K, Centeno TP, Vidal RO, Rahman R-U, Benito E, Fischer A, Dennerlein S, Rehling P, Feussner I, Bonn S, Simons M, Urlaub H, Rizzoli SO. 2018. Precisely measured protein lifetimes in the mouse brain reveal differences across tissues and subcellular fractions. Nat Commun 9(1):4230.

Gigant B, Curmi PA, Martin-Barbey C, Charbaut E, Lachkar S, Lebeau L, Siavoshian S, Sobel A, Knossow M. 2000. The 4 Å X-Ray Structure of a Tubulin:Stathmin-like Domain Complex. Cell 102(6):809–816.

Gilbert Di Paolo, Robert Lutjens, Véronique Pellier, Stephen A. Stimpson, Marie-Hélène Beuchat, Stefan Catsicas, Gabriele Grenningloh. 1997. Targeting of SCG10 to the Area of the Golgi Complex Is Mediated by Its NH2-terminal Region. J Biol Chem 272:5175–5182.

Grenningloh G, Soehrman S, Bondallaz P, Ruchti E, Cadas H. 2004. Role of the microtubule destabilizing proteins SCG10 and stathmin in neuronal growth. J Neurobiol 58(1):60–69.

Guerra San Juan I, Nash LA, Smith KS, Leyton-Jaimes MF, Qian M, Klim JR, Limone F, Dorr AB, Couto A, Pintacuda G, Joseph BJ, Whisenant DE, Noble C, Melnik V, Potter D, Holmes A, Burberry A, Verhage M, Eggan K. 2022. Loss of mouse Stmn2 function causes motor neuropathy. Neuron 110(10):1671-1688.e6.

Himi T, Okazaki T, Wang H, McNeill TH, Mori N. 1994. Differential localization of SCG10 and p19/stathmin messenger RNAs in adult rat brain indicates distinct roles for these growth-associated proteins. Neuroscience 60(4):907–926.

Honnappa S, Cutting B, Jahnke W, Seelig J, Steinmetz MO. 2003. Thermodynamics of the Op18/Stathmin-Tubulin Interaction*. Journal of Biological Chemistry 278(40):38926–38934.

Hsu PD, Scott DA, Weinstein JA, Ran FA, Konermann S, Agarwala V, Li Y, Fine EJ, Wu X, Shalem O, Cradick TJ, Marraffini LA, Bao G, Zhang F. 2013. DNA targeting specificity of RNA-guided Cas9 nucleases. Nat Biotechnol 31(9):827–832.

Hung V, Zou P, Rhee H-W, Udeshi ND, Cracan V, Svinkina T, Carr SA, Mootha VK, Ting AY. 2014. Proteomic Mapping of the Human Mitochondrial Intermembrane Space in Live Cells via Ratiometric APEX Tagging. Molecular Cell 55(2):332–341.

Jourdain L, Curmi P, Sobel A, Pantaloni D, Carlier M-F. 1997. Stathmin: A Tubulin-Sequestering Protein Which Forms a Ternary T 2 S Complex with Two Tubulin Molecules. Biochemistry 36(36):10817–10821.

Kalocsay M. 2019. APEX Peroxidase-Catalyzed Proximity Labeling and Multiplexed Quantitative Proteomics. In: Sunbul M, Jäschke A, editors. Proximity Labeling: Methods and Protocols. New York, NY:Springer. p 41–55.

Klim JR, Williams LA, Limone F, Guerra San Juan I, Davis-Dusenbery BN, Mordes DA, Burberry A, Steinbaugh MJ, Gamage KK, Kirchner R, Moccia R, Cassel SH, Chen K, Wainger BJ, Woolf CJ, Eggan K. 2019. ALS-implicated protein TDP-43 sustains levels of STMN2, a mediator of motor neuron growth and repair. Nat Neurosci 22(2):167–179.

Krus KL, Strickland A, Yamada Y, Devault L, Schmidt RE, Bloom AJ, Milbrandt J, DiAntonio A. 2022. Loss of Stathmin-2, a hallmark of TDP-43-associated ALS, causes motor neuropathy. Cell Rep 39(13):111001.

Labun K, Montague TG, Krause M, Torres Cleuren YN, Tjeldnes H, Valen E. 2019. CHOPCHOP v3: expanding the CRISPR web toolbox beyond genome editing. Nucleic Acids Research 47(W1):W171–W174.

Leonard DG, Ziff EB, Greene LA. 1987. Identification and characterization of mRNAs regulated by nerve growth factor in PC12 cells. Mol Cell Biol 7(9):3156–3167.

Lutjens R, Igarashi M, Pellier V, Blasey H, Paolo GD, Ruchti E, Pfulg C, Staple JK, Catsicas S, Grenningloh G. 2000. Localization and targeting of SCG10 to the trans-Golgi apparatus and growth cone vesicles. European Journal of Neuroscience 12(7):2224–2234.

Marklund U, Larsson N, Gradin HM, Brattsand G, Gullberg M. 1996. Oncoprotein 18 is a phosphorylation-responsive regulator of microtubule dynamics. The EMBO Journal 15(19):5290–5298.

Melamed Z, Lopez-Erauskin J, Baughn MW, Zhang O, Drenner K, Sun Y, Freyermuth F, McMahon MA, Beccari MS, Artates JW, Ohkubo T, Rodriguez M, Lin N, Wu D, Bennett CF, Rigo F, Da Cruz S, Ravits J, Lagier-Tourenne C, Cleveland DW. 2019. Premature polyadenylation-mediated loss of stathmin-2 is a hallmark of TDP-43-dependent neurodegeneration. Nat Neurosci 22(2):180–190.

Mori N, Morii H. 2002. SCG10-related neuronal growth-associated proteins in neural development, plasticity, degeneration, and aging. Journal of Neuroscience Research 70(3):264–273.

Niethammer P, Bastiaens P, Karsenti E. 2004. Stathmin-Tubulin Interaction Gradients in Motile and Mitotic Cells. Science 303(5665):1862–1866.

Pellier−Monnin V, Astic L, Bichet S, Riederer BM, Grenningloh G. 2001. Expression of SCG10 and stathmin proteins in the rat olfactory system during development and axonal regeneration. Journal of Comparative Neurology 433(2):239–254.

Poulain FE, Sobel A. 2007. The “SCG10-LIke Protein” SCLIP is a novel regulator of axonal branching in hippocampal neurons, unlike SCG10. Molecular and Cellular Neuroscience 34(2):137–146.

Prudencio M, Humphrey J, Pickles S, Brown A-L, Hill SE, Kachergus JM, Shi J, Heckman MG, Spiegel MR, Cook C, Song Y, Yue M, Daughrity LM, Carlomagno Y, Jansen-West K, Castro CF de, DeTure M, Koga S, Wang Y-C, Sivakumar P, Bodo C, Candalija A, Talbot K, Selvaraj BT, Burr K, Chandran S, Newcombe J, Lashley T, Hubbard I, Catalano D, Kim D, Propp N, Fennessey S, Fagegaltier D, Phatnani H, Secrier M, Fisher EMC, Oskarsson B, Blitterswijk M van, Rademakers R, Graff-Radford NR, Boeve BF, Knopman DS, Petersen RC, Josephs KA, Thompson EA, Raj T, Ward M, Dickson DW, Gendron TF, Fratta P, Petrucelli L. 2020. Truncated stathmin-2 is a marker of TDP-43 pathology in frontotemporal dementia. J Clin Invest 130(11).

Ravelli RBG, Gigant B, Curmi PA, Jourdain I, Lachkar S, Sobel A, Knossow M. 2004. Insight into tubulin regulation from a complex with colchicine and a stathmin-like domain. Nature 428(6979):198–202.

Rhee H-W, Zou P, Udeshi ND, Martell JD, Mootha VK, Carr SA, Ting AY. 2013. Proteomic Mapping of Mitochondria in Living Cells via Spatially Restricted Enzymatic Tagging. Science 339(6125):1328–1331.

Riederer BM, Pellier V, Antonsson B, Di Paolo G, Stimpson SA, Lütjens R, Catsicas S, Grenningloh G. 1997. Regulation of microtubule dynamics by the neuronal growth-associated protein SCG10. PNAS 94(2):741–745.

Shin JE, Geisler S, DiAntonio A. 2014. Dynamic regulation of SCG10 in regenerating axons after injury. Experimental Neurology 252:1–11.

Shin JE, Miller BR, Babetto E, Cho Y, Sasaki Y, Qayum S, Russler EV, Cavalli V, Milbrandt J, DiAntonio A. 2012. SCG10 is a JNK target in the axonal degeneration pathway. PNAS 109(52):E3696–E3705.

Steinmetz MO, Jahnke W, Towbin H, García−Echeverría C, Voshol H, Müller D, Oostrum J van. 2001. Phosphorylation disrupts the central helix in Op18/stathmin and suppresses binding to tubulin. EMBO reports 2(6):505–510.

Steinmetz MO, Kammerer RA, Jahnke W, Goldie KN, Lustig A, Oostrum J van. 2000. Op18/stathmin caps a kinked protofilament-like tubulin tetramer. The EMBO Journal 19(4):572–580.

Summers DW, Milbrandt J, DiAntonio A. 2018. Palmitoylation enables MAPK-dependent proteostasis of axon survival factors. PNAS 115(37):E8746–E8754.

Tararuk T, Östman N, Li W, Björkblom B, Padzik A, Zdrojewska J, Hongisto V, Herdegen T, Konopka W, Courtney MJ, Coffey ET. 2006. JNK1 phosphorylation of SCG10 determines microtubule dynamics and axodendritic length. J Cell Biol 173(2):265–277.

Thornburg-Suresh EJC, Richardson JE, Summers DW. 2023. The Stathmin-2 membrane-targeting domain is required for axon protection and regulated degradation by DLK signaling. J Biol Chem 299(7):104861.

Westerlund N, Zdrojewska J, Padzik A, Komulainen E, Björkblom B, Rannikko E, Tararuk T, Garcia-Frigola C, Sandholm J, Nguyen L, Kallunki T, Courtney MJ, Coffey ET. 2011. Phosphorylation of SCG10/stathmin-2 determines multipolar stage exit and neuronal migration rate. Nature Neuroscience 14(3):305–313.

