## Supplementary Figures for "Tubulin Regulates the Stability and Localization of STMN2 by Binding Preferentially to Its Soluble Form"

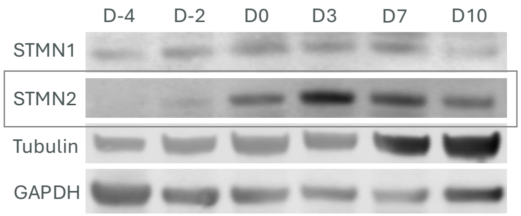


**Supplementary Fig. 1.** The expression of STMN2 peaked on the third day after plating.

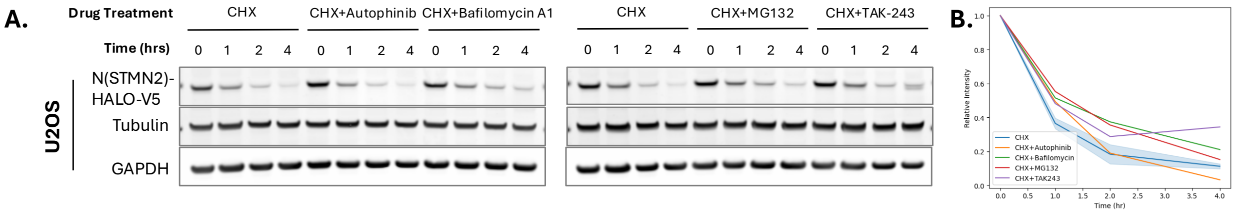


**Supplementary Fig. 2.**

**A.** Western blot of CHX chase assay along with 10 μM Autophinib, 200 nM Bafilomycin A1, 5 μM MG132, or 200 nM TAK-243 in N(STMN2)-HALO-V5 expressing U2OS cells. The CHX-only condition samples shown on the left and right panels are from the same experiment and were loaded twice as controls for the western blot.

**B.** Relative band intensities of N(STMN2)-HALO-V5 (E), plotted against chase time. For each condition, band intensities are normalized to the initial time point (0 hr). Data points for the two CHX-only conditions are plotted with confidence intervals.

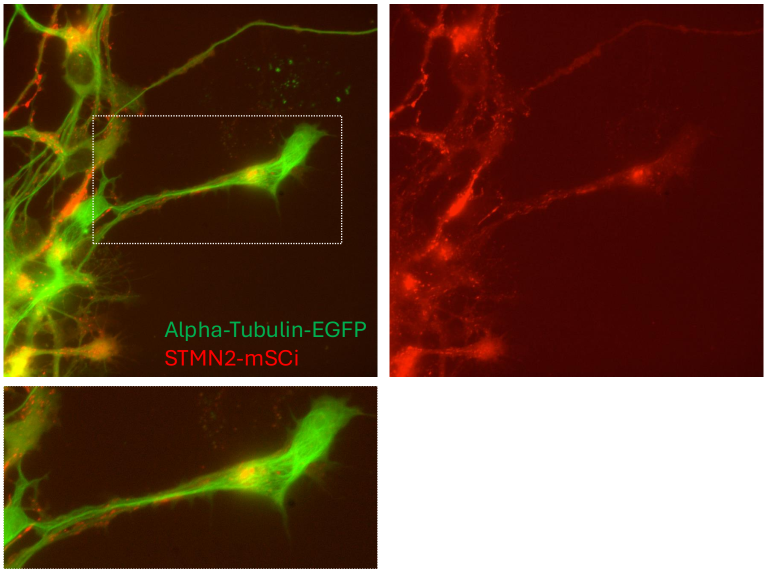


**Supplementary Fig. 3.** D3 iNs expressing TUBA-EGFP and STMN2-mSCi showed no obvious colocalization.
